# Florida *Drosophila melanogaster* genomes sampled 13 years apart show increases in warm-associated SNP alleles

**DOI:** 10.1101/2020.10.23.352732

**Authors:** Krishna R. Veeramah, Evgeny Brud, Walter F. Eanes

## Abstract

We studied genetic change in *Drosophila melanogaster* using whole-genome SNP data from samples taken 13 years apart in Homestead, FL. This population is at the southern tip of a well-studied US latitudinal cline. On the non-inversion-carrying chromosome arms, 11-16% of SNPs show significant frequency changes. These are enriched for latitudinal clines and genic sites. For clinal SNPs each allele is either the northern- or southern-favored. Seventy-eight to 95 percent with significant frequency increase are southern-favored. Five to seven percent of SNPs also show significant seasonal change and involve increases in the northern-favored allele during the season. On the 2L and 3R chromosome arms there are significant seasonal shifts for common inversions. We identify regions and genes that are candidates for selection. These regions also show correspondence with those associated with soft sweeps in Raleigh, NC. This shift towards southern-favored alleles may be caused by climate shifts or increased African-European admixture.

## INTRODUCTION

Understanding genetic change in present-day populations is important because of (1) specific interests in how natural populations respond to local climate change (Anderson *et al*. 2005; Cogni *et al*. 2017; Nadeau *et al*. 2017; Umina *et al*. 2005), and (2) the broader question of the general intensity of contemporary selection (Hancock 2016; Hansen *et al*. 2012; Messer *et al*. 2016). We address this question in *Drosophila melanogaster* by examining genome-wide change in a population from south Florida (FL) using samples collected in 1997 and then again 13 years later. The Homestead, FL locality sits at the southern end of a well-characterized latitudinal cline on the east coast of the US (Bergland *et al*. 2014; Fabian *et al*. 2012; Sezgin *et al*. 2004). The cline runs from northern temperate to southern subtropical regions, presenting populations with markedly different environmental challenges. Along with a comparable latitudinal cline in eastern Australia, this iconic eastern North American cline has featured in more than 70 published studies over the last four decades (though mostly in the last 15 years), either as the primary focus or providing the context of studying specific genes or traits of interest (Adrion *et al*. 2015). While both Australian and US clines may have arisen as the result of a migration-selection process after European invasion, there is evidence that ongoing or past European-African admixture may be involved in originating the cline for many genes (Bergland *et al*. 2015; Kao *et al*. 2015b; Kaplan & Weir 1995; Yukilevich & True 2008a; Yukilevich *et al*. 2010). In this regard, the long-term genetic stability of the Florida population in the face of potential migration from African-Caribbean sources is certainly relevant to evaluating the evolutionary response to modern-day selection pressures.

Bergland et al. (2014) collected whole-genome data across the North American cline to identify SNPs that exhibited regular seasonal oscillations in a temperate orchard in eastern Pennsylvania (PA). Their study used Pool-seq methods to estimate allele frequencies and also assessed latitudinal changes in the same SNPs in six populations evenly spaced from Maine to Florida. The PA site, in the upper temperate section of the cline, was sampled in the spring and fall in each of three successive years (2009-2011). They identified a set of ~1,750 SNPs that under their criteria appeared to show regular seasonal variation. Moreover, population differentiation in SNPs measured as F_ST_ was found to be a predictor of those SNPs that also showed significant seasonal cycling.

In 1997, a collection of *D. melanogaster* isofemale lines was made by Brian Verrelli in Homestead, Florida at the Fruit and Spice Park (botanical garden). This collection was part of a study to look at latitudinal clines in a small number of specific enzyme polymorphisms (Sezgin *et al*. 2004; Verrelli & Eanes 2001). In 2008 and 2010, three collections were also made from the same park as part of Bergland et al. (2014). These later collections have been used in part to study the diapause associated *cpo* gene (Cogni *et al*. 2015; Cogni *et al*. 2014; Schmidt *et al*. 2008).

Integrating the 1997 and 2008/2010 samples provides a unique opportunity to look at differences that may have emerged in this southern end of the North American cline over the past decade. Therefore we subjected the 1997 sample to Pool-seq and combined it with the extensive Pool-seq data from Bergland et al.(2014). This combination allows us to quantify allele frequency SNP differences for two collections that are separated by 13 years or seasons, and approximately 190 *D. melanogaster* generations. By identifying those SNPs possessing both significant latitudinal clines and significant frequency change, we can place the incidence of 13 year change in the context of any systematic bias supporting either the southern- or northern-favored allele of each SNP.

## RESULTS

We generated Pool-seq resequencing data from 60 pooled isofemale lines collected from Homestead, Florida in 1997 (henceforth designated HFL97). Mean coverage when mapped against genome build dm5 using BWA was 423x. For comparative purposes we reprocessed Pool-seq data from all 13 clinal populations from Bergland et al. (2014) using the same pipeline. Mean coverage in these samples ranged from 29-207X. In order to avoid potential ascertainment biases resulting from the higher coverage of our HFL97 sample, we limit all our subsequent analyses to SNPs originally identified in Bergland et al. (2014), with additional filtering based on coverage ranges (to eliminate potential copy number variants, CNVs) and the estimated minor allele frequency being >10% in all populations, resulting in a final set of 768,302 SNPs across all five chromosome arms.

### General Distribution of pairwise F_ST_ values

Estimation of genome-wide mean pairwise F_ST_ amongst all 14 populations demonstrated that HFL97 was genetically most similar to the two populations from Bergland et al.(2014) also collected from Homestead, FL1 and FL2. In general, genetic similarity decreases with distance from Homestead (Supplemental Figure 1). These results suggest that the Homestead population in 2010 is largely the same population as that sampled 1997, rather than being a recent immigrant population from elsewhere. Therefore, in subsequent analyses we focused on comparisons amongst the three Homestead populations HFL97, FL1 and FL2. FL1 is designated a “spring” collection based on pooled sampling from Homestead in July 2008 and July 2010, while FL2 is designated a “fall” collection based on sampling in December 2010. Table 1 presents the summary F_ST_ statistics based on a block bootstrap sampling of 10 kb intervals across each arm. We note that HFL97 is genetically more similar to FL2 than FL1, consistent with their same season of collection. At a genome-wide level (ignoring the inversion-polymorphic arms 2L and 3R, which show generally more unpredictable patterns, see below), the F_ST_ between HFL97 and FL2 is similar in magnitude to the F_ST_ between FL1 and FL2 (Table 1), demonstrating the strong effect of seasonal selection on SNPs and associated frequency shifts at linked sites, despite FL1 and FL2 being so close temporally. However, the F_ST_ between HFL97 and FL1 is noticeably larger than between FL2 and FL1, consistent with additional long-term change over the separation of ~190 generations or 13 seasonal cycles. Therefore, to evaluate the contribution of seasonal changes within the Homestead population, we compared the Bergland spring and fall collections (FL1 versus FL2), and to investigate long-term change we compared the 1997 and 2010 December collections (HFL97 versus FL2).

**Table 1.** Summary F_ST_ Statistics

| Table 1.<br>Summary<br>F<sub>ST</sub><br>Statistics<br>Arm | Comparison | $F_{ST} \times 100$ | 5-95% CI |
| --- | --- | --- | --- |
| 2R | HFL97 v FL2 | 1.32 | 1.25-1.32 |
|  | HFL97 v FL1 | 1.83 | 1.76-1.83 |
|  | FL1 v FL2 | 1.36 | 1.29-1.36 |
| 2L | HFL97 v FL2 | 1.12 | 1.07-1.12 |
|  | HFL97 v FL1 | 2.07 | 1.99-2.07 |
|  | FL1 v FL2 | 1.56 | 1.48-1.56 |
| 3R | HFL97 v FL2 | 2.08 | 1.98-2.08 |
|  | HFL97 v FL1 | 2.05 | 1.97-2.05 |
|  | FL1 v FL2 | 1.17 | 1.11-1.17 |
| 3L | HFL97 v FL2 | 1.24 | 1.19-1.24 |
|  | HFL97 v FL1 | 1.71 | 1.65-1.71 |
|  | FL1 v FL2 | 1.17 | 1.11-1.17 |
| X | HFL97 v FL2 | 1.56 | 1.43-1.56 |
|  | HFL97 v FL1 | 2.41 | 2.27-2.41 |
|  | FL1 v FL2 | 1.47 | 1.32-1.47 |

### Seasonal shift detected between the FL1 and FL2 collections

Bergland et al. (2014) identified a subset of approximately 1750 SNPs in the Pennsylvania population that showed significant evidence of seasonal oscillation in allele frequency. In Pennsylvania, the temperature averages between winter and summer seasons are in the range of ~8C to 26C. However, the difference in Homestead, Florida is less extreme in the winter, averaging 20C in July and 27C in December. While we lack the repetitive sampling of Pennsylvania over three consecutive years, the impact of seasonal change in Florida nevertheless can be examined by comparing the FL1 and FL2 samples. This further allows us to interpret the 13 year change within the context of seasonality. We focus first on the chromosome arms 2R, 3L and the X chromosome, and address the major inversion-carrying arms separately.

We find that 4.9%, 7.2%, and 6.4% of SNPs on the X, 2R, and 3L arms demonstrate a significant frequency difference between the FL1 and FL2 collections (*P* < 0.05). Amongst this subset there was a significant enrichment for SNPs that have been identified as showing latitudinal clines (log_2_ = 0.129, *P* < 2.3 × 10^-7^). There is also a significant enrichment for synonymous SNPs (log_2_= 0.171, *P* < 1.16 × 10^-9^) (non-synonymous SNPs were enriched in a similar direction but their smaller total numbers likely explains the non-significance); there is significant under-enrichment of intergenic SNPs (log_2_ = −0.143, *P* < 5.28 × 10^-9^; Figure 1A). Figure 2A shows the SNP frequency changes from spring (FL1) to late fall (FL2) for the southern-favored allele (those identified as being clinal in Bergland et al. with *q* < 0.10) for each SNP with significant frequency change (*P* < 0.05) against its position.

**Figure 1.**
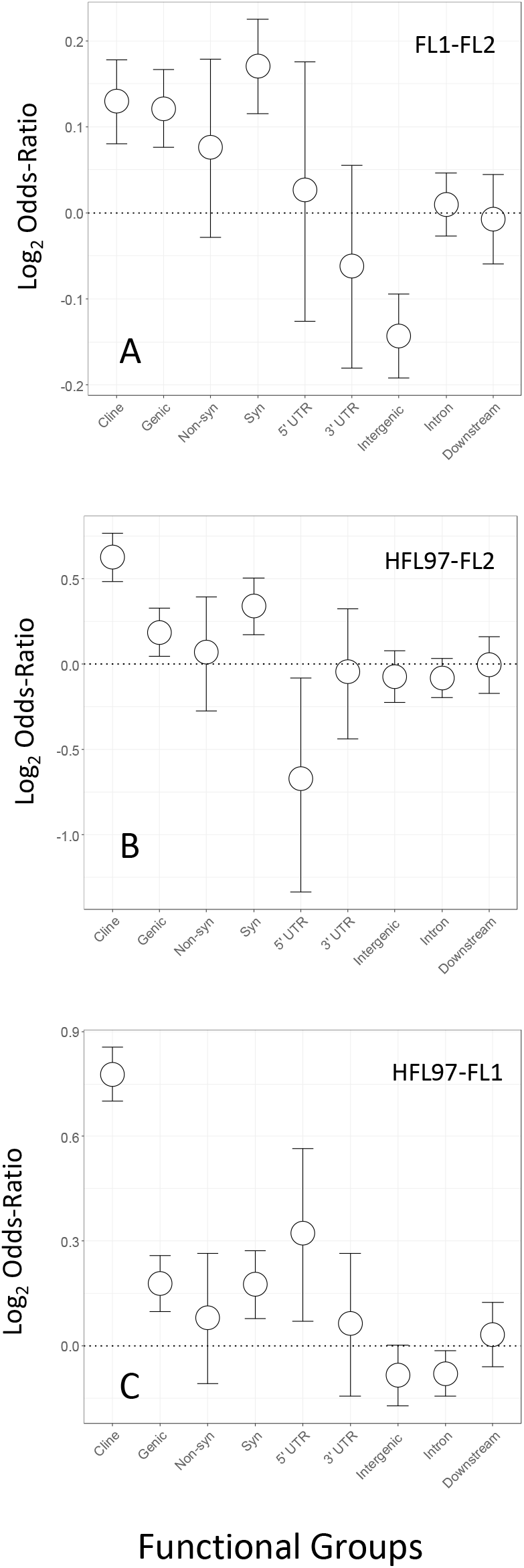
SNP enrichment (as log_2_ odds ratios) for clines and different functional classes. Arms X, 2R and 3L combined. (A)Seasonal (FL1-FL2) collections. (B) Thirteen year interval December collections (HFL97-FL2) for SNPs with *q* < 0.10. (C) Thirteen year interval July-December collections (HFL97-FL1), for SNPs with *q* < 0.10.

**Figure 2.**
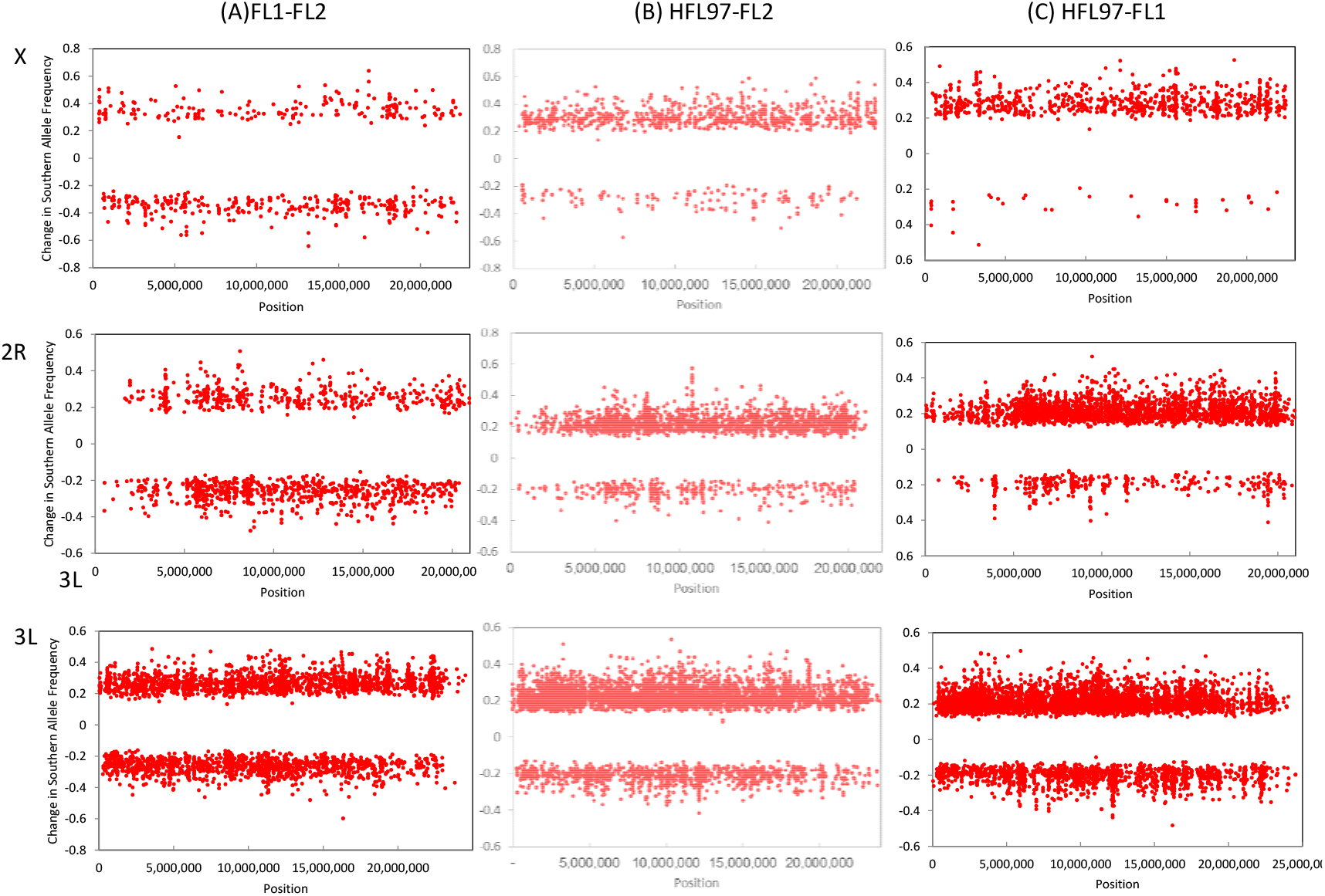
Southern-favored allele frequency changes (A) FL1-FL2 that are both significant at *P* < 0.05 and clinal at *P* < 0.05 plotted against chromosome position for the X chromosome, 2R, and 3L arms. (B) HFL97-FL2 that are significant at *P* < 0.05 and that are also clinal against chromosome position for the X chromosome (clinal at *P* < 0.05), 2R (*q* < 0.10), and 3L (*q* < 0.10) arms. (C) HFL97-FL1 that are significant at *P* < 0.05 and that are clinal against chromosome position for the X chromosome (clinal at *P* < 0.05), 2R (*q* < 0.10) and 3L (*q* < 0.10) arms.

The confidence in assessing north-south bias in frequency change depends on our confidence in defining clinal SNPs. Overall, the proportion of SNPs possessing significant clines (at *P* < 0.05) varies among chromosomes (Bergland *et al*. 2014). The X chromosome possesses a much lower proportion of clinal SNPs (14% for *P* < 0.05), while for chromosome arm 2R, 9% of the SNPs show clines and the incidence for 3L was much higher at 27%. Figure 3A shows for each arm the proportion of SNPs where the southern alleles are significantly increasing (*P* < 0.05), when defining clinal SNPs using varying FDRs from *q* < 0.20 to 0.005. For the X chromosome, we find that, depending on clinal *q*-value cutoff, between 66 to 90% of the SNP alleles increasing from July to December are northern-favored. For the 2R arm, sample sizes are limited, but we see that about 63% of the seasonally-changing SNPs are northern-favored. The opposite is observed for the 3L arm where the direction of this enrichment is for southern-favored alleles.

**Figure 3.**
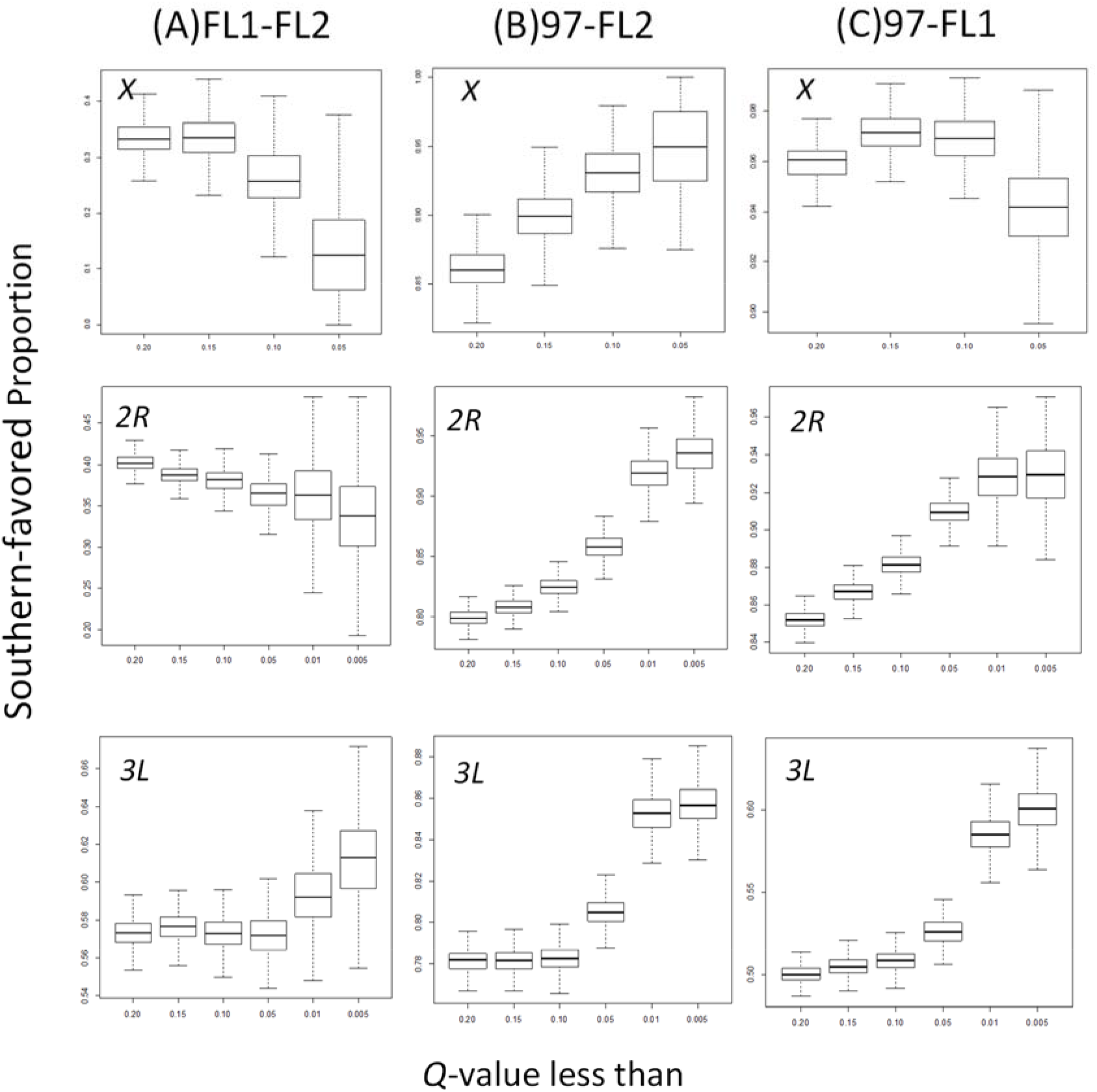
Box plots of all X, 2R and 3L SNPs with significant shifts in allele frequency (*P* < 0.05) and divided into FDR subsets with different clinal *q*-values. Each boxplots represents 1,000 bootstrap sample mean estimates. (A) FL1-FL2 collections (B) HFL97-FL2 collections. (C) HFL97-FL1 collections.

There was no overlap between our SNPs showing significant frequency differences on the X and a set of seasonally varying SNPs previously identified using seasonal data from Pennsylvania, partly because the latter are highly underrepresented in the Bergland et al.(2014) data (only 55 from a genome-wide total of 1,756 SNP using an FDR of *q* < 0.3). For chromosome arm 2R, there were only 7 SNPs with significant differences between FL1 and FL2 that overlapped with the 362 seasonal SNPs from Bergland et al. (2014) for which the frequency change from spring to fall was in the same direction (*P* < 0.349, P-value obtained by bootstrapping a random set of control SNPs 1,000 times). Interestingly 11 SNPs overlapped where the frequency change was in the opposite direction, a result that was significant (*P* < 0.036). For chromosome arm 3L, we see the opposite pattern, with 13 SNPs overlapping 496 known seasonal SNPs in the same direction (*P* < 0.068), and 8 in the opposite direction (*P* < 0.563). Overall there appears to be a only minor directional shift associated with either latitudinal or seasonal selection between FL1 and FL2, and the signals are diametrically opposed between chromosomes.

### Major enrichment of southern-favored alleles from December 1997 to December 2010

We next examined evidence for systematic allele frequency changes over a 13-year period related to the clinal and seasonal SNPs identified in Bergland et al (2014). Given the observation of a real (though subtle) seasonal component to SNP changes in Florida, we consider changes between the December 1997 (HFL97) and the two collections in 2008/2010 (FL1 and FL2) separately. As with the seasonal comparison we see evidence for enrichment of clinal SNPs (the set with *q* < 0.10) among SNPs that significantly shift in frequency (*P* < 0.05) over the HFL97-FL2 time frame (Figure 1B; log_2_ = 0.627, *P* < 2.2 × 10^-16^). There is a clear significant enrichment for synonymous SNPs in the pooled arms (Figure 1B; log_2_ = 0.341, *P* < 6.79 × 10^-5^) (non-synonymous are again the same direction, but not significant) and a marginal under-enrichment of 5’UTR SNPs (log_2_= −0.67, *P* < 0.023). In the comparison of chromosome arms X, 2R, 3L, we observed that 7.4, 10.4, and 9.9% of SNPs possess a significant frequency difference (P < 0.05) (Figure 2B).

For all three arms, the clear majority of SNPs with significant change (*P* < 0.05) and significant clines involve increases in the southern-favored allele (Figure 3B). When conditioning those with significant frequency differences and then varying clinal SNP sets with decreasing *q*-values (increasing our confidence in clinal assessment), the proportion of increasing alleles that are also southern-favored increases to about 95% of the SNPs on all three inversion-free arms (Figure 3B).

### Major enrichment of southern-favored alleles December from 1997 to July 2008/2010

Finally, we might expect a contrast of differences between December 1997 and July 2008/2010 when combining both seasonal and 13-year changes, as suggested in the earlier F_ST_ analysis. We observe that 8.8, 13.4, and 12.4% of SNPs on these chromosome arms possess a significant frequency difference (*P* < 0.05). There is a clear enrichment for clinal (log_2_=0.66, *P* < 2.2 × 10^-6^), synonymous (log_2_ = 0.198, *P* < 9.73 × 10^-5^) and 5’UTR (log_2_ = 0.318, *P* < 0.012) SNPs in the joint set with a difference FDR of *q* < 0.10 (Figure 1C). In contrast, intron SNPs are marginally under-enriched (log_2_ = −0.080, *P* < 0.016), as well as intergenic SNPs (log_2_ = −0.085, *P* < 5×10^-2^). The large majority of SNPs also show increases in southern-favored alleles (Figures 2C, Figure 3C).

When conditioning those SNPs with significant frequency differences (*P* < 5%) and varying clinal SNP sets with decreasing *q*-values, the proportion of increasing alleles that are southern-favored increases to about 94% on both the X and 2R arms (Figure 3C). The increase on 3L is much smaller and presumably reflects the counter selection for northern alleles on this arm (Figure 3C).

### The major inversion bearing chromosome arms 2L and 3R

The Homestead population segregates for inversions on all four autosomal arms, but the large *In(2L)t* and *In(3R)Payne* inversions each segregate at frequencies of ~25%. Both *In(2R)NS* and *In(3L)P* are relatively rare in the Homestead population and the inversions *In(3R)Mo, In(3R)C*, and *In(3R)K* are virtually absent (Kapun et al. 2016a; Kapun & Flatt 2019; Mettler et al. 1977; Sezgin et al. 2004). Both long-term and seasonal variation in SNP frequencies across the 2L and 3R chromosome arms are expected to be strongly affected by frequency changes in the *In(2L)t* and *In(3R)Payne* inversions. This is evident in the plots of SNP frequency change against position (Figures 4 and 5). Since, the frequencies of both inversions are well known to decrease with increasing latitude (Kapun et al. 2016a; Mettler et al. 1977; Sezgin et al. 2004), we define them as southern-favored arrangements.

**Figure 4.**
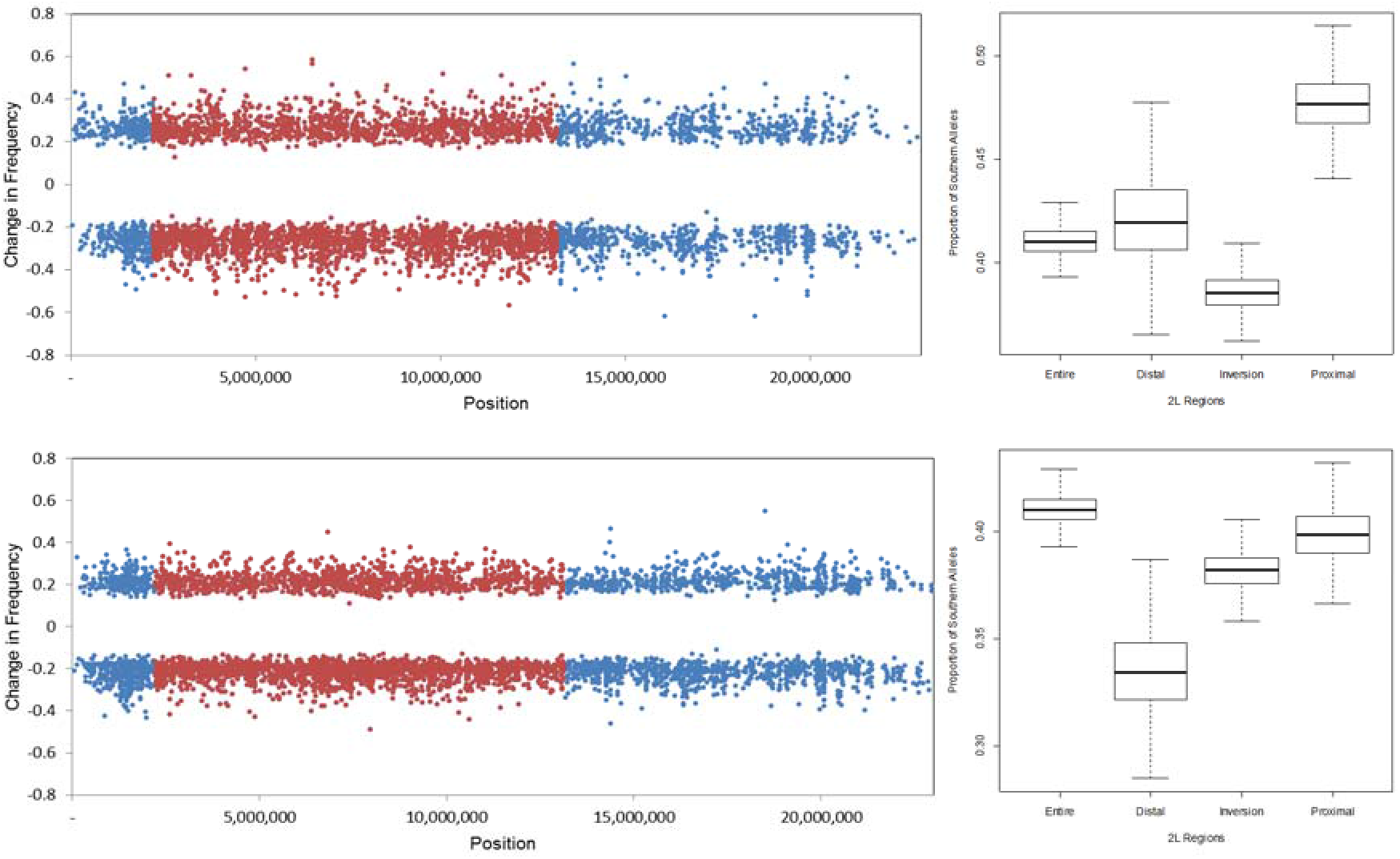
Plot of all SNPs with both significant clines (*q* < 0.10) and significant change in frequency of southern alleles (*P* < 0.05) plotted against position on arm 2L. Red points are SNPs within the *In(2L)t* inversion. (Top) Seasonal data FL1-FL2. (Bottom) Long-term HFL97-FL2 data. Inserts to right are boxplots represents 1,000 bootstrap sample mean estimates of southern-favored allele proportions for SNP changes (*P* 0.05) and clines (*q* < 0.10) for positions inside and outside the inversions.

**Figure 5.**
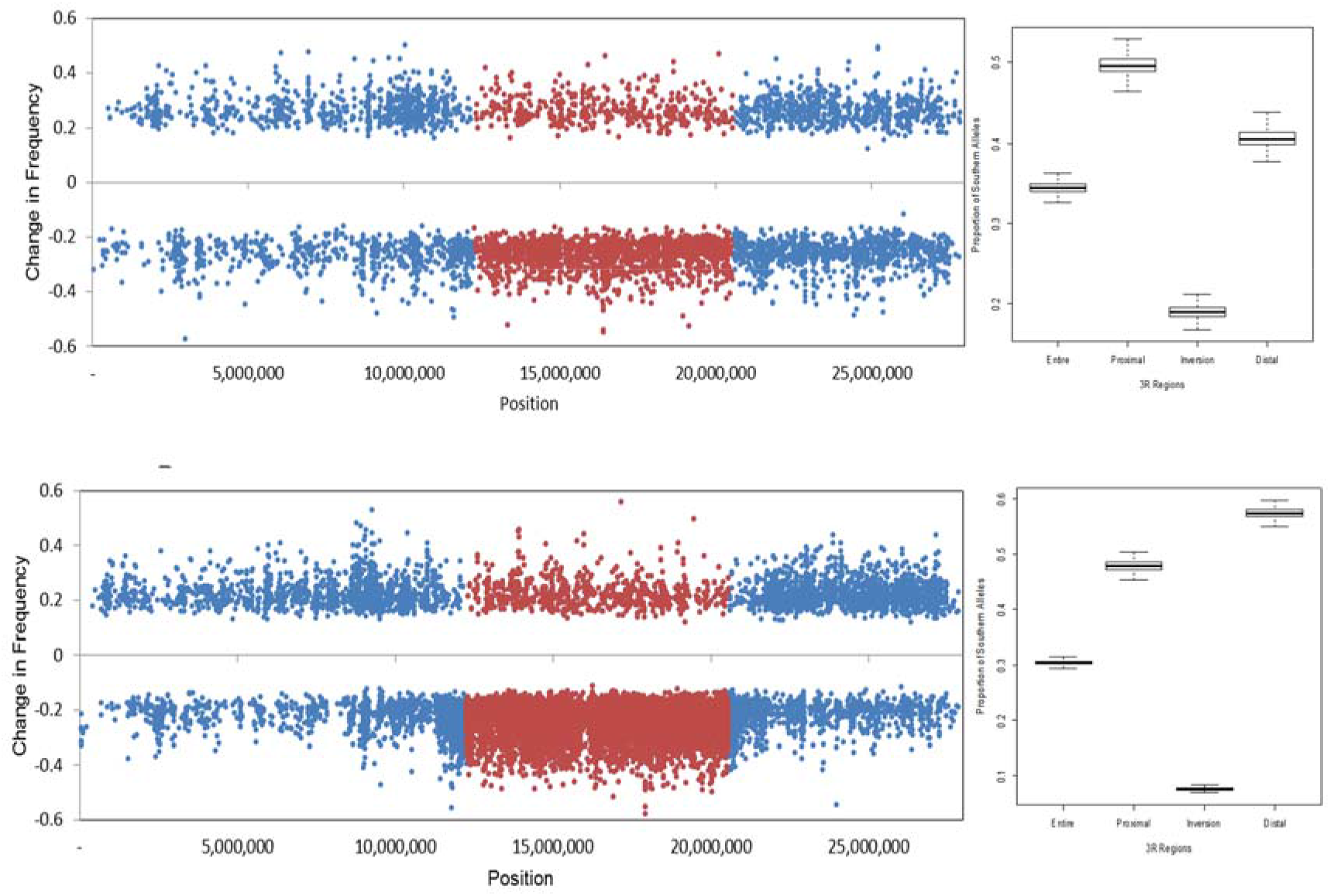
Plot of all SNPs with both significant clines (*q* < 0.10) and significant change in frequency of southern alleles (*P* <0.05) plotted against position on arm 3R. Red points are SNPs within the *In(3R)Payne* inversion. (Top) Seasonal FL1-FL2. (Bottom) Long-term HFL97-FL2 data. Inserts to right are boxplots represents 1,000 bootstrap sample mean estimates of southern-favored allele proportions for SNP changes (*P*< 0.05) and clines (*q* < 0.10) for positions inside and outside the inversions.

Diagnostic markers near the breakpoints have been identified for the *In(2L)t* and *In(3R)Payne* inversions (Kapun *et al*. 2016a; Kapun *et al*. 2014), although several of these diagnostic SNPs are not segregating in our samples. Averaging the frequencies of the six diagnostic SNPs reported for *In(2L)t*, we see there is an apparent seasonal (FL1 to FL2) drop of 23% in the frequency of this inversion (Kapun *et al*. 2014). In contrast, there is *no significant difference* in the frequency of *In(2L)t* (~3%) between the December collections in 1997 and 2010 (*P* < 0.34 across six diagnostic SNPs). This inversion therefore exhibits notable seasonal change, yet its frequency is the same between the 13-year collections.

In contrast, using a single diagnostic SNP for *In(3R)Payne* there is estimated to be a marginally significant long-term drop in *In(3R)Payne* frequency of 25% (P < 0.043). There is also an apparent 12 percent seasonal drop between the July 2008/2010 (FL1) and December 2010 (FL2) samples, but this is not significant (*P* < 0.27, *q* < 0.79). For the most part, these observations are strengthened by the study of the changes of internal SNPs with respect to northern- and southern-favored status.

When looking at the northern-southern allele status of SNPs that show significant frequency change, we need to first note that the inversion clines in *In(2L)t* and *In(3R)Payne apparently* drive the high percentage of SNPs that are clinal on these arms (29% and 44% on 2L and 3R respectively). Figure 5 (top) shows the distribution across the entire 2L arm of southern-favored allele frequency changes for the FL1 to FL2 seasonal period. For the entire arm, a large proportion of the subset of SNPs with significant changes (*P* < 0.05) are also clinal and for the clinal subset with *q* < 0.10 we observe that 59% involve spring to fall increases in the northern-favored allele in each SNP. This depends weakly on SNP location. For SNPs *within* the *In(2L)t* inversion, we observe that 61 percent involve increases in the northern-favored allele. Though this general pattern extends into the flanking regions outside the inversion, the signal is more subtle although the bootstrap estimates do not overlap.

For arm 2L, Figure 4 (bottom) shows the southern-favored allele frequency changes from HFL97 to FL2. Again, using *P* < 0.05 for frequency differences and *q* < 0.10 as a FDR cutoff for clines, we observe for the entire arm a small, but nonetheless significant excess (62%) of the northern alleles are increasing in frequency. The flanking proximal and distal regions also possess small but nonsignificant shifts favoring northern-favored alleles. The general observation is that there is an apparent seasonal change in *In(2L)t*, but we observed overall long-term stasis in its frequency. This is also predicted by the diagnostic SNPs.

Across the 3R arm, we see that 6.4% of the SNPs differ significantly between the seasonal FL1 and FL2 collections (*P* < 0.05) and this involves 65% of the northern-favored alleles increasing (Figure 5 top). For SNPs *within* the inversion 81% of the northern-favored alleles are increasing in frequency with season. The small distal telomeric region shows subtle shifts of increasing northern-favored alleles (60%), while there is no notable pattern favoring either northern or southern alleles in the proximal region towards the centromere.

We see that from HFL97 to FL2 across the entire 3R arm, about 70% of the alleles that significantly (*P* < 0.05) vary in frequency are northern-favored (Figure 5 bottom). Within the *In(3R)Payne* inversion we observe that 92% of the increasing alleles are northern-favored. In contrast to the dramatic shift seen within the inversion, the distal and proximal flanking region exhibit small shifts in the opposite direction favoring southern alleles and this increases with distance from the inversion breakpoints, and therefore these regions respond as the majority of the genome. Overall, we conclude that in this population at the extreme southern end of the latitudinal cline there has been a 13-year decrease in the *In(3R)Payne* inversion. Again, both the seasonal and 13-year changes mirror the predictions from using the inversion-diagnostic SNP changes for both arms.

### F_ST_ Outliers and Correspondence with Other Studies

Mean F_ST_ values for the 2R arm are 0.0196, 0.0148, and 0.0156 for HFL97-FL1, HFL97-FL2, and FL1-FL2 data sets, respectively. Figure 6 (A-C) plots the mean of a modified version of the *population branch statistic* (*PBS*) computed from pairwise *F*_ST_ estimates amongst HFL97 and FL2 (Shriver *et al*. 2004; Yi *et al*. 2010), for 5 kb windows sliding every 1 kb along the X, 2R and 3L arms, with the *PBS* just for the long-term changes (Figure 6A-C) (see Methods and Materials).

**Figure 6.**
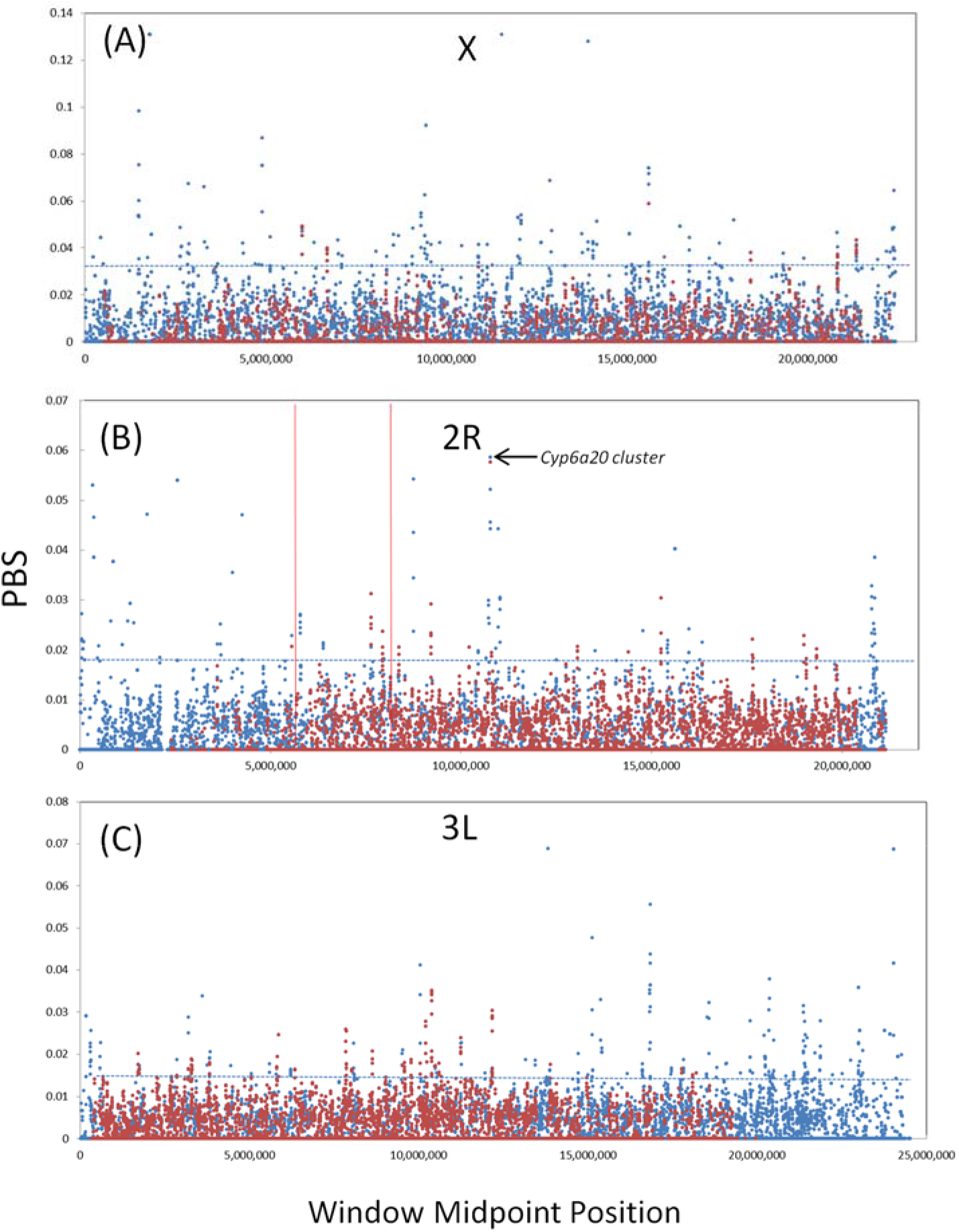
HFL97-FL2 sliding plot of mean PBS values for 5 kb windows and sliding increments of 1 kb against mean midpoint position (blue points). The dotted line represents the cutoff for all windows in the upper 1%. Windows containing of *n* > 23, 36 and 38 SNPs for the X and 2R and 3L arms are highlighted in red and genes found in these windows are listed in Supplemental Tables 1-3 for each arm. The positions of candidate soft sweep regions from Garud et al. (2015) that overlap with the top 1% windows are shown with red vertical lines.

The number of SNPs varies across the 5 kb windows reflecting the well-known genome-wide relationship between genetic diversity and local recombination (Langley *et al*. 2012; Mackay *et al*. 2012). Because the variance in mean *F*_ST_ among windows strongly depends on the number of SNPs in each window, in the final analysis we have included only windows with greater than 26, 36, and 38 SNPs for the X, 2R and 3L arms, respectively. This removes windows with spuriously high *F*_ST_ estimates simply because they possess small numbers of SNPs. Supplemental Tables 1-3 list, after the joining of the contiguous windows in the top 1% of the most extreme *PBS* values for each arm, our candidate windows and the individual genes contained in each. This cutoff identified 18, 37, 48 windows potentially responding to long-term selection for these arms (X, 2R, 3L), respectively.

While a number of interesting genes are found in these windows, the highest ranked region on the 2R arm is a 5 kb window of 41 SNPs with a *PBS_LT_* value of 0.057 (percentile < 10^-4^). This window contains three genes, *Cyp6a9, Cyp6a19*, and *Cyp6a20*. The SNP in this window with the highest *PBS* value (PBS = 0.208 and pairwise *F*_ST_ between HFL97 and FL2 = 0.495) involves an amino acid change at position 10,770,502 (G289D) in the *Cyp6a20* gene where the minor allele, G, has increased from 0.336 in HFL97 to 0.745 and 0.898 in the FL1 and FL2 samples respectively. By coalescing several flanking windows (albeit using slightly lower SNP numbers) this window can be expanded to create a 10 kb window with high *PBS* estimates (0.0442-0.0558), and includes the gene, *Cyp6a23*. This entire region of 10 kb contains eight cytochrome P450 genes. The window has a depth of coverage for each pool consistent with the average of the genome, so unlike the previously described case of the cytochrome P450 gene *Cyp6g1*, it is not a duplicated region (Schmidt *et al*. 2010). The entire SNP set in this block is represented by well-supported clines (*q* < 0.01) with southern-favored alleles increasing between the 13-year interval collections.

We were interested in the correspondence of our significant *PBS* windows with the Garud et al. (2015) study that identified regions of selection soft-sweeps in the 2005 Raleigh, NC (DGRP) autosomal genome sequences. Again we studied just the inversion-free autosomal arms. On the 2R arm they identify 13 soft sweep regions varying in size from 11kb to 764kb. Finally, in our long-term analyses of 2R, two of our top 0.1% (39 regions) significant *PBS* windows fall in the 13 candidate soft sweep regions. In the first, we find five contiguous *PBS* windows spanning 9kb containing three genes (*CG34234, fdl, DH44-R2*) fall within their large (764kb) soft sweep region that contains 121 genes. A second *PBS* window, centered at 5,562,500, and containing only the *Mmp2* gene is found in the their soft-sweep window spanning 216kb that carries 23 genes. For the 3L arm neither of the two regions identified as targets of soft sweeps in Raleigh correspond to windows of high *PBS* (top 0.1%) in our data.

## DISCUSSION

In this study we examine systematic genetic change in a subtropical *D. melanogaster* population by studying two collections from Homestead, Florida separated by 13 years. To do this, we have taken isofemale samples collected in 1997 that have been stored in ethanol in an ultracold freezer for 19 years, and used the Pool-seq approach to compare this sample to the identical locality collected in 2008-2010 by Bergland et al. (2014). The 13-year time span between samples reflects about 190 generations at this latitude (Pool 2015). Several prominent features emerge from the joint analysis of these data sets.

### Inversion-limited Arms X, 2R and 3L

We should first note that the SNPs showing significant shifts in both the seasonal and long-term time scales are significantly enriched for clinal and genic (especially synonymous) SNPs. The most prominent observation is that for those SNPs with significant increasing frequency and also with statistically significant latitudinal clines, the southern-favored allele at each SNP constitutes the clear majority of alleles increasing in frequency between the 13-year interval samples. The pattern becomes most apparent as the FDR applied to the significance of clines becomes more stringent and increases our confidence in each of the north-south assignment of alleles at each SNP. It is important to emphasize that this pattern of apparent long-term increase in southern-favored alleles takes place in a more subtle seasonal context of increases in northern-favored alleles from the spring to fall. Comparing the two FL1 and FL2 samples that were collected in July (2008/2010) and December (2010), we see that the majority of the allele frequency shifts on the X and the 2R arm from spring to fall involve the *decrease* of southern-favored alleles, while curiously the opposite is observed on the 3L arm. The reason for this difference between arms is unclear, but it should be noted that the frequency of SNPs with significant clines identified by Bergland et al. (2014) also differs markedly between arms and are more abundant on the 3L arm.

Without year-to-year replication of collections we cannot tell if the observed spring-fall frequency changes in Florida reflect a regular seasonal oscillation or are just the general level of variation expected between two collections at different times in the same year. Bergland et al. (2014) collected their Pennsylvania orchard samples over three years with the goal of recovering a set of statistically well-supported seasonally oscillating SNPs and looking at their collective properties. They also asked if those oscillating alleles favored early in the summer (spring collections) and conversely favored in the fall were also southern and northern-biased across the cline. They first observed that spatially varying SNPs were more likely to be oscillating temporally, similar to the cline enrichments reported here, however their results suggest that spring (July) collections in PA favored more northern-associated alleles, while fall collections (October-December) were more southern-favored. This is the opposite of what we observe in this southern FL population. Their study was from a temperate seasonal population found in the upper portion of the latitudinal cline, so this may reflect fundamental differences in how selection is acting across the season in the southern subtropical and northern temperate regions, but nonetheless it suggests that in general the same selection forces acting across the latitudinal cline are partially acting in a seasonal fashion as well.

### Major Inversion-bearing Arms

For the chromosomal arms 2L and 3R, interpreting any pattern of favoring the northern and southern-favored status of alleles must take into account the cosmopolitan *In(2L)t* and *In(3R)Payne* inversions. These segregate at appreciable frequencies in Florida and are themselves clinal and decreasing northward. Recognizing this, we can discern a couple of generalities about both inversions and discuss them in the context of earlier studies and reviews (Kapun *et al*. 2016b; Kapun & Flatt 2019). First, there is observed to be a notable seasonal shift with the *In(2L)t* inversion decreasing in frequency by about 23% from July to December. This is consistent with the observation that *In(2L)t* seasonally cycles in a temperate Linvilla, PA orchard (Bergland *et al*. 2014; Kapun *et al*. 2016b). Interestingly, the direction of seasonal frequency change of this inversion depends on the spatial context along the cline, with the inversion increasing into the late season at the southern end of the cline and decreasing into late season in the northern population. Second, the frequency of *In(2L)t* appears to have not changed, averaging about 25% in the two December collections sampled 13 years apart.

In contrast, the overall data for *In(3R)Payne* predict a lower frequency of the inversion in this sample 13 years later. This is of note given the earlier report of dramatic *increase* in the frequency of *In(3R)Payne* over 20 years in Australia that is interpreted to be the result of climate change (Umina *et al*. 2005). *In(3R)Payne* also shows a seasonal shift, dropping in frequency consistent with the Homestead population in the Kapun et al. (2016a) (see their Supplemental Table S1), but again in the opposite direction to the seasonal change observed in Linvilla, PA (Bergland *et al*. 2014; Kapun *et al*. 2016b). Note, the Kapun et al. (2016a) study also used the Sezgin et al (2004) data so some concordance across studies is expected.

Congruence between our high PBS (or F_ST_) windows and earlier reported candidate regions for soft selection are important because they help discover genes likely to be recurrent targets of selection in different populations. For the 2R and 3L arms, we find a few examples of overlap between regions identified in our *PBS* outlier scan and the regions of soft selection proposed by Garud et al. (Garud *et al*. 2015; Garud & Petrov 2016) from their analysis of the Raleigh, NC (DGRP) genomes. For example, the region they discovered around the insecticide-resistance gene *Cyp6g1* (Schmidt *et al*. 2010) continues to also hold interest in our data. This soft sweep candidate region is broad (764 kb and 121 genes) and overlaps our individual long-term outlier window that contains just the *CG34234, fdl*, and *DH44-R2* genes. Overall, this simply says that these apparent soft sweep regions identified in the Raleigh population do correspond to hot spots of long-term selection in the south FL population. This is significant because the convergent identification of these sites comes from both different data and populations, and using different methods. The soft sweeps were discovered using extended haplotype statistics (Sabeti *et al*. 2002) in the Raleigh, NC population, while our study is based on observed frequency change in a long-term study in a different population at the tip of the cline. In our FL data, the most obvious *PBS* peak spans a large cluster of six cytochrome P450 genes. The most prominent gene with the highest F_ST_ SNPs is *Cyp6a20*, which uniquely has been identified independently as associated with aggression behavior (Wang & Anderson 2009; Wang et al. 2008).

What are the possible reasons for the increase in southern-biased alleles between the 13-year collections? One cannot rule out a response to climate change, where the south FL population is adapting to a warmer environment. An alternative explanation would be demographic, involving admixture from an African source: this is a question of increasing interest in studies of these clines (Bergland *et al*. 2015; Flatt 2016) in both North America and Australia. Recent studies, two from the DGRP collection (sampled in Raleigh, NC, in 2005) (Duchen *et al*. 2013; Pool 2015) and one from a widespread sample of 17 whole genomes collected from the SE US and Caribbean islands (Kao *et al*. 2015b) have predicted admixture between European and African sources. Overall, admixture in the DGRP genomes is estimated at about 19% African, but notably with little evidence of admixture on the X chromosome (Pool 2015). African admixture in the genomes from the SE US and Caribbean islands is predicted to vary from near zero to 50%, but generally increases towards the Caribbean (Kao et al. 2015b). Generally, the X chromosome appears to be exceptional with respect to admixture and a number of other features. Pool et al. (2012) observed that European admixture into African populations was underrepresented on the X chromosome. F_ST_ estimates between Europe and sub-Saharan Africa show greater differentiation for genes on the X (Kao *et al*. 2015b). Our observation that, compared to the two autosomal arms, the X chromosome shows the greater proportion of alleles that are increasing and southern-favored under long-term change points to the X chromosome as fundamentally different.

If admixture between European and African sources is ongoing in the southeastern US, then an outcome might include chromosome-dependent hybrid incompatibilities and the occurence of adaptive introgression (Hedrick 2013). There are reports of mating and fitness incompatibilities between crosses from lines in this zone of admixture (Kao *et al*. 2015a; Yukilevich & True 2008b; Yukilevich & True 2008c). These incompatibilities might be expected to have different associations with the X and autosomes because of both sex-dependent rates of introgression and of the expectations of Haldane’s Rule that genetic incompatibilities will arise more frequently on the X chromosomes (Charlesworth *et al*. 1987). Two other observations are also relevant to this question. The discovery by Pool et al. (2012) of European admixture in some African populations also found large variation in European-African ancestry among genomes within localities. This large individual variation suggests local departures from random mating that is associated with European ancestry. Moreover, in these same African populations the X chromosome shows much less European admixture. This same disparity between the X and autosomes is observed for admixture in the Raleigh population (Pool 2015) while we see in our study that the increase in southern-favored alleles between 1997 and 2010 collections is most prominent on the X chromosome. A potential explanation for this is that while Haldane’s Rule acting on X-linked deleterious alleles may limit the overall amount of introgression on the X chromosome compared to the autosomes (as is also seen in other systems, include Neanderthal introgression into humans [Harris and Neilsen 2016]), adaptive alleles acquired from the introgression populations may also experience stronger positive selection due to the faster-X effect (Charlesworth et al.2018).

The original interest in this cline began with the assumption that it is a migration-selection cline (Oakeshott *et al*. 1983; Oakeshott *et al*. 1981; Oakeshott *et al*. 1982). That assumption must now include demographic history (Flatt 2016). This does not rule out selection-migration as playing a causal role in the clinal variation in many genes, but future studies must include the additional possibility of ongoing African admixture and adaptive introgression. This does not rule out that much of the clinal variation reflects selection acting on the original variation brought to the US from temperate European populations, an estimated 150 years ago (Keller 2007). Nonetheless, this is now confounded by African introductions from the Caribbean that are also segregating for the same original polymorphisms that were introduced into Europe much earlier. Teasing apart these conflicting phenomena will require a detailed in-depth study using full genome sequences across the cline.

## Methods and Materials

### Pool-seq of HFL97

The HFL97 collection was made by Brian Verrelli in December 1997 at Fruit and Spice Park and is listed in Verrelli and Eanes (2001). Isofemale lines were kept in the lab for extraction of second and third chromosomes, and then each line was eventually stored for 19 years at −70C in 100% ethanol. DNA was extracted from 60 pooled isofemale lines (using two males/line), with yields from single flies ranging from 0.15 to 0.69 *ug*. From the pooled males 5.2 ug of DNA was prepared and submitted to the New York Genome Center (NYGC) for library preparation and sequencing. The lines showed some degradation with a bimodal size distribution with peaks at ~300bp and ~2,500 bp. The NYGC used a PreCR Repair Mix to perform fragment repair as a precautionary measure. Size selection was then performed to separate the two major peaks, and the 2,500 bp fragments only were sheared. Both the unsheared and sheared fragments were then used for library preparation before sequencing on a single lane of an Illumina HiSeq 2500 1TB upgrade to generate paired-end 2 x 125bp reads. Following the processing described below, the 1997 HFL average coverage was 425X with 10% duplicate rate and 96% of unique reads mapping to the v5.39 *D. melanogaster* (dm5) reference genome.

### Bioinformatic Processing

Samples underwent an pipeline where reads were first trimmed using the software Skewer (Jiang *et al*. 2014) and then mapped to *dm5* using bwa-mem (Li & Durbin 2009). Duplicate reads were identified and marked using Picard Tools (http://broadinstitute.github.io/picard) and indels were realigned using GATK Indel Realigner (DePristo *et al*. 2011). Finally, called bases underwent base quality recalibration using GATK BQSR to produce a finished BAM file for further analysis. SNPs identified in DGRP freeze 2 were used to recalibrate base qualities. We applied this pipeline to both our preliminary data and the original raw Bergland et al. (2014) Pool-seq read data.

### Allele Frequency Estimation

Allele frequencies were estimated at all 1,119,372 SNPs identified in Bergland et al. (2014) using the maximum-likelihood estimator of Lynch et al. (2014) (https://github.com/kveeramah/Lynch-PoolSeq-estimator). We implemented the estimator as described in the original paper, including an initial pass to estimate sequencing error, *ε*, from allele frequencies *p* < 0.90 and then re-estimating all SNPs with *p* > 0.90 given this *ε* (though ultimately we ignored all SNPs with a minor allele frequency less than *p* < 0.10). We filtered against mapping and base phred-scaled qualities less than 20 before calling SNPs and discarded sites with less than a third or greater than twice the mean coverage to guard against duplicate regions (i.e. CNVs). Though estimated using slightly different methods, the allele frequencies estimated in our pipeline and the original estimates from Bergland et al. (2014) were very similar, with a mean *r*^2^ = 0.97 (min = 0.94, max = 0.99).

We used the likelihood ratio tests of Lynch et al. (2014) to test for significant frequency differences between any two populations. To first build a set of SNPs with significant change in frequencies, *α* < 5% was used, and more subsequent cutoff (e.g. *q* < 10%) were used to build joint sets, but these were constrained by reductions in sample size. These are discussed in the text, case by case.

### F_ST_

Estimation of pairwise F_ST_ was performed using Popoolation2 (Kofler *et al*. 2011) using the approach described in Karlsson et al. (2007). This estimator takes into account chromosome number in the pool. The Bergland et al. (2014) samples are composed entirely of individual male flies (except for North Carolina, which contained females) collected on site and pooled for Pool-seq and thus the chromosome counts are straightforward. In contrast, the 1997 samples are sampled from individual isofemale lines that were maintained in the lab for short period between being frozen and stored. Since each line is descended from a single mated female collected in the wild, two males are sampled from each isofemale line, so the effective number of chromosomes is 240 autosomes and 120 X chromosomes. Again, we ignored mapping and base qualities less than 20.

### Modified PBS

Schriver et al. (2004) previously described the estimation of locus-specific branch lengths based on pairwise F_ST_ estimates amongst three populations. Yi et al.(2010) further developed this into the Population Branch Statistic (PBS) via a log-transformation that could be used to identify in the presence of an outgroup which of a pair of populations was likely under selection due to an elevated *F_ST_* at a particular locus. Here we adapt the PBS approach in order to identify loci amongst our three Florida pool-seq populations that may be under (a) long term and (b) seasonal selection. We first define the three pairwise F_ST_S, *F_ST1_* = HFL97 v FL1, *F_ST2_* = HFL97 v FL2, and *F_ST3_* = FL1 v FL2. Under no selection, drift and gene flow we expect these values to be equal, though we would expect *F_ST1_* and *F_ST2_* to be bigger than *F_ST3_* due to drift over the 13 years (Supplemental Figure 2A). If there has been strong long-term selection at a locus from 1997 and 2010 that is distinct from any seasonal selection, we might expect both *F_ST1_* and *F_ST2_* to be large and *F_ST3_* to be small (Supplemental Figure 2B). Thus we define:

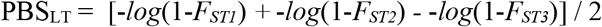

It is possible to have a large PBS_LT_ if either *F_ST1_* is exceptionally large but *F_ST2_* is not (and vice-a-versa), but this would not be consistent with long term selection. Therefore, we also ignore loci (functionally setting the value to zero) where |*F_ST1_ − F_ST2_*| > |*F_ST3_ - F_ST1_*| or |*F_ST3_ - F_ST2_*|.

Similarly, if there has been strong seasonal cycling selection that is distinct from any long-term selection, then we expect *F_ST1_* and *F_ST3_* to be large and *F_ST2_* to be small (Supplemental Figure 2C). Thus we define:

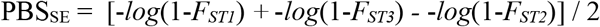

Again, it is possible to have a large PBS_SE_ if either *F_ST1_* is large but *F_ST3_* is not (and vice-a-versa), but this would not be consistent with seasonally-cycling selection. Therefore we also ignore loci (functionally setting the value to zero) where |*F_ST1_ - F_ST3_*| > |*F_ST1_ - F_ST2_*| or |*F_ST2_ - F_ST3_*|.

In both cases we looked for the largest PBS statistics as evidence of selection. We apply our modified PBS on *F_ST_*S calculated in 5kb windows sliding every 1kb for each chromosome separately. We identified all windows in the 99% tail of the chromosome, joining any significant windows in which the nearest boundary points were separated by no more than 1kb (for example, if there were two significant window spanning positions 10,000-15,000 and 16,000-21,000, they were joined). We then performed the same analysis on individual SNPs within these significant regions ±250kb.

### Assigning alleles as northern or southern-favored

A consideration in our analysis is the systematic shifting of the overall characteristic of the SNPs that is associated with the environment. For each biallelic SNP polymorphism that possesses a statistically significant latitudinal cline at the applied FDR, it is possible to assign to the reference and alternate alleles the status of either “southern-” or “northern-favored” depending respectively on a negative or positive allele frequency association with increasing latitude. The confidence in that assignment depends on the statistical support for the associated cline from the Bergland et al. (2014) data. The *P* and *q* values for the clinal significance of each SNP are taken directly from that data set. For SNPs with significant GLM regressions, the sign of the correlation was used to assign status for the reference allele. A reference allele that increases in frequency with increasing latitude is classified as northern-favored and vice versa. A FDR cutoff *q* < 10% for each cline test was used to assemble a high stringency assignment of northern or southern favored status for 2R and 3L. For the X chromosome to assemble reasonable sample sizes a cutoff of *q* < 27% was applied.

## Acknowledgements

The authors acknowledge National Institutes of Health Grant, GM-090094 to WFE.

## SUPPLEMENTAL FILES

**Supplemental Figure 1.**
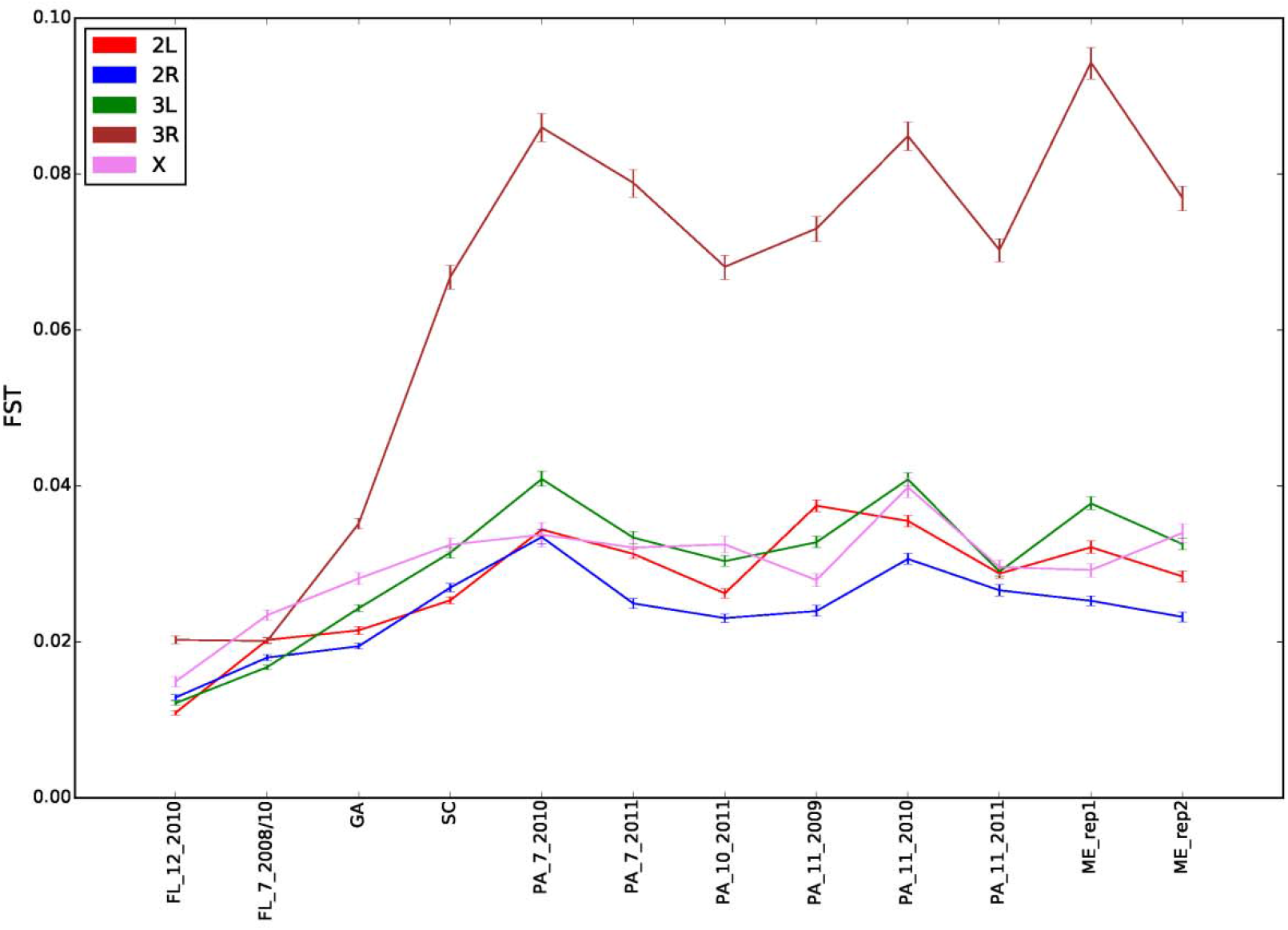
Plot of average chromosome pairwise F_ST_ of Bergland et al. pool-seq samples (excluding North Carolina because of residual heterozygosity and unclear temporal sampling) against HFL97. CIs generated using 1000 10kb block bootstrapping.

**Supplemental Figure 2.**
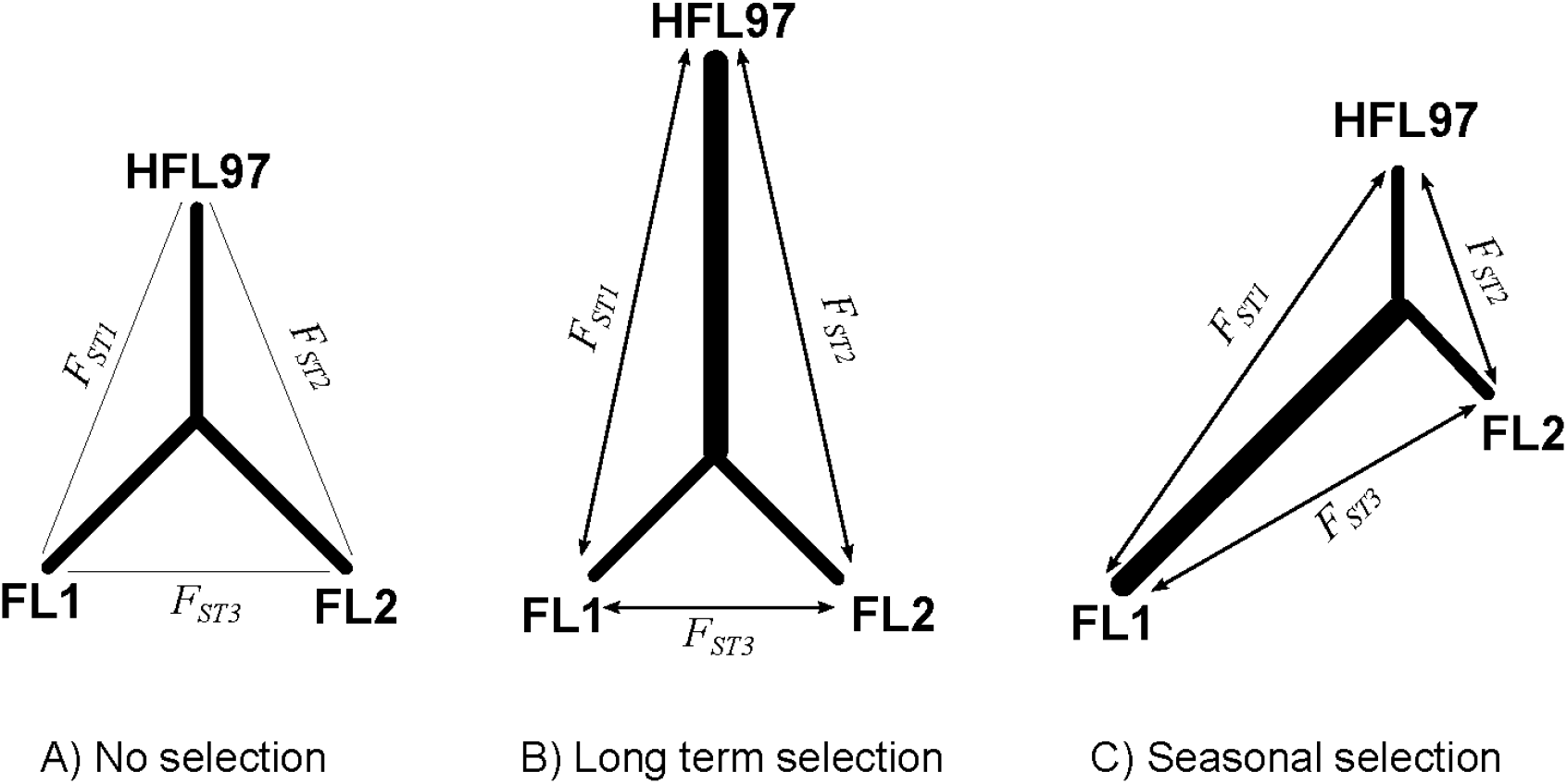
Expected relative FSTs under different selection regimes

